# Male violence disrupts estrogen receptor β signaling in the female hippocampus

**DOI:** 10.1101/2023.09.23.559092

**Authors:** Jacopo Agrimi, Lucia Bernardele, Naeem Sbaiti, Marta Canato, Ivan Marchionni, Christian U. Oeing, Beatrice Vignoli, Marco Canossa, Nina Kaludercic, Claudia Lodovichi, Marco Dal Maschio, Nazareno Paolocci

**Affiliations:** Department of Biomedical Sciences, University of Padova, Padova, Italy; Department of Medicine, Johns Hopkins University School of Medicine, Baltimore, MD, USA; Department of Internal Medicine and Cardiology, Charité University Medicine, Berlin, Germany; Department of Cellular, Computational, and Integrative Biology, University of Trento, Trento, Italy; Neuroscience Institute -CNR Padova, Italy; Veneto Institute of Molecular Medicine, Padova, Italy

**Keywords:** Intimate Partner Violence, Domestic Violence, Estrogen Receptor, BDNF, TrkB, Corticosterone

## Abstract

Women are the main target of intimate partner violence (IPV), which is escalating worldwide. Mechanisms subtending IPV-related disorders, such as anxiety, depression and PTSD, remain unclear. We employed a mouse model molded on an IPV scenario (male *vs.* female prolonged violent interaction) to unearth the neuroendocrine alterations triggered by an aggressive male mouse on the female murine brain. Experimental IPV (EIPV) prompted marked anxiety-like behavior in young female mice, coincident with high circulating/cerebral corticosterone levels. The hippocampus of EIPV-inflicted female animals displayed neuronal loss, reduced BrdU-DCX-positive nuclei, decreased mature DCX-positive cells, and diminished dendritic arborization level in the dentate gyrus (DG), features denoting impaired neurogenesis and neuronal differentiation. These hallmarks were associated with marked down-regulation of estrogen receptor β (ERβ) density in the hippocampus, especially in the DG and dependent prosurvival ERK signaling. Conversely, ERα expression was unchanged. After EIPV, the DG harbored lowered local BDNF pools, diminished TrkB phosphorylation, and elevated glucocorticoid receptor phosphorylation. In unison, ERβ KO mice had heightened anxiety-like behavior and curtailed BDNF levels at baseline, despite enhanced circulating estradiol levels, while dying prematurely during EIPV. Thus, reiterated male-to-female violence jeopardizes hippocampal homeostasis in the female brain, perturbing ERβ/BDNF signaling, thus instigating anxiety and chronic stress.

## Main

“Intimate partner violence” (IPV) is the physical/sexual violence, stalking, or psychological injury perpetrated by a current or former partner/spouse^1^. IPV can endanger the function of multiple organs and has an undeniable and appalling feature: the victims are primarily women^2^. IPV neurological and psychiatric repercussions have monopolized the lens of scientists and caregivers, especially epidemiologically^3^. Yet how IPV ignites/exacerbates brain/behavioral disorders, especially when reiterated, remains to be deciphered in full. Experimental models replicating human IPV’s critical features are crucial to addressing this unknown. To fill this gap, we adapted a recently validated animal model that leads a female mouse to social defeat after a violent, prolonged interaction with an aggressive male (Experimental IPV, EIPV)^4^. Using fertile female mice, we tested whether EIPV downregulates estrogen-mediated signaling, focusing on the hippocampus, a primary hub of mood control and a well-documented target of acute/chronic stress of different etiology^5^. Within it, we zoomed on the dentate gyrus (DG), a prominent neurogenesis site in the adult brain^6^. More in detail, we tested whether, at the hippocampal/DG level, EIPV alters estrogen receptor (ER) expression patterns and dependent prosurvival ERK signaling that, in turn, controls the expression of brain-derived neurotrophic factor (BDNF)^7^. BDNF is essential for proper neuronal development and response to stress conditions of different etiology in adulthood^8^. We grounded our hypothesis on evidence showing that ERβ selective agonists, particularly, exert potent anxiolytic effects during animal behavioral testing by inhibiting the ACTH and corticosterone response to stress^9^. Moreover, estrogen can shape BDNF expression in many ways. First, by convergence, when interacting with BDNF receptors, namely tyrosine kinase receptor B (TrkB): estrogen and BDNF/TrkB activate transcription factors, such as CREB, leading to the transcription of a battery of prosurvival genes, including *bdnf* itself^7^. Or by induction. Indeed, BDNF contains a canonical estrogen response element, and estrogen can also directly modify the BDNF promoter epigenetically^7^.

### EIPV triggers anxiety-like behavior in female mice

We tailored a recently validated social defeat protocol in female mice^4^ borrowed from the resident-intruder paradigm. The latter is widely used to investigate human psychosocial stress through animal models^10^. In doing so, as shown in Fig.1A, we reproduced primary features of human IPV, such as the reiteration of physical contact/violence and psychological threat.

**Fig. 1.**
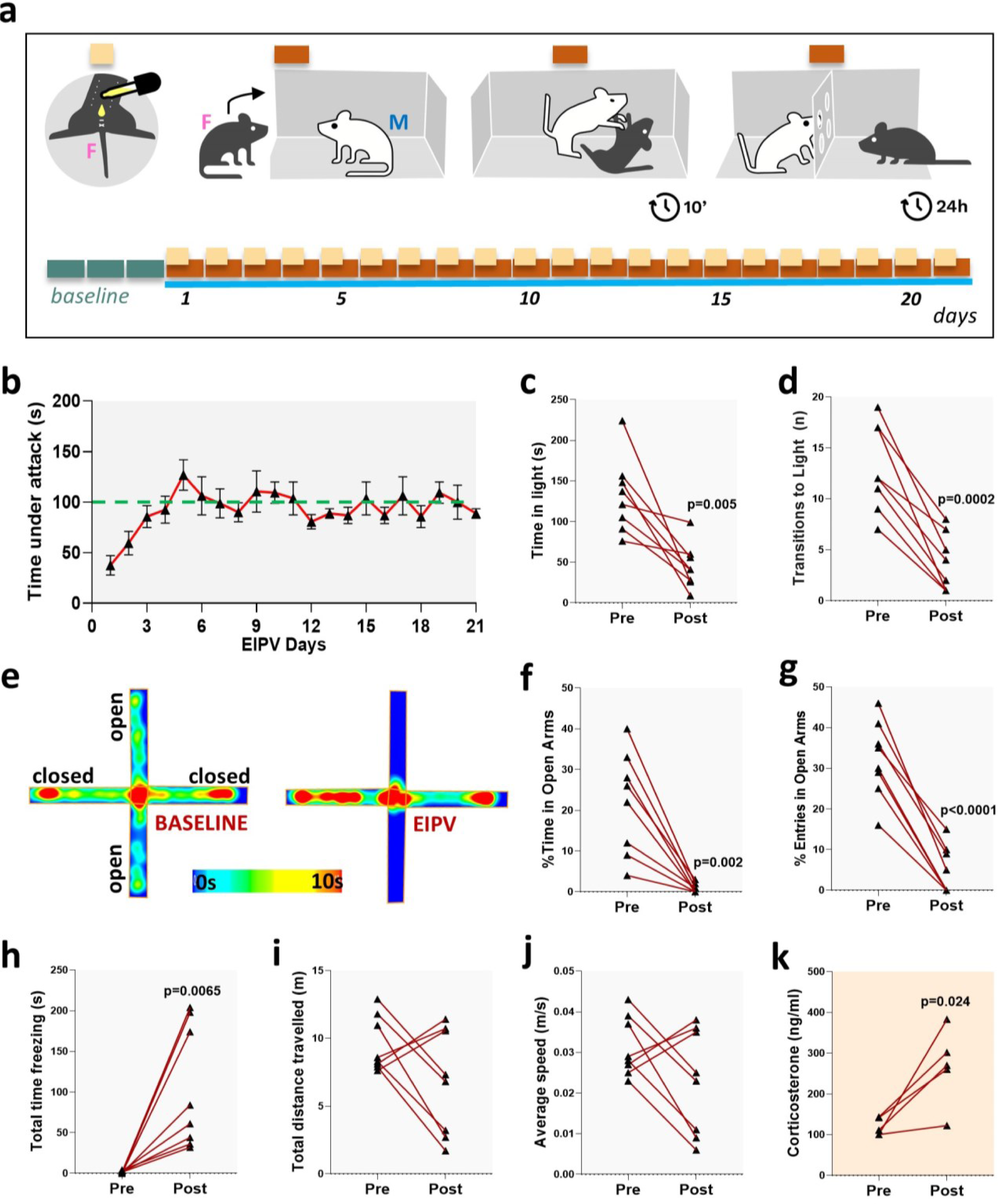
ERβ and EIPV alter BDNF levels in the DG. **a**, Representative images of IF staining for BDNF in the DG (hilar cells), WT, and ERβ KO mice before and after EIPV. **b,** Quantification of IF staining for BDNF in the DG, control WTs vs. ERβ KO mice, before and after EIPV, n=5, Kruskal–Wallis one-way test.

Consistently, the EIPV procedure led to daily physical male-toward-female attacks (computed in ≅100s per day) (Fig.1B) while mixing this physical, aggressive behavior with a sensorial intimation on a recurrent basis. Following 21 days of this treatment, we set out to determine whether the planned EIPV procedure was penetrant enough to perturb the female mouse’s behavior, focusing on exploratory activity and anxiety. Therefore, at the end of the procedure, the female mice were subjected to behavioral tests, including the light-dark box (LDB) and the elevated plus maze (EPM)^11^. Then, we compared post-EIPV female mice’s behavior with baseline performance and assayed the pre- and post-EIPV levels of corticosterone, the primary stress hormone in rodents^12^. The two main parameters of the LDB test revealed a significant surge of anxiety-like behavior coupled with reduced exploratory activity in female mice after EIPV. More in detail, they spent significantly less time in the light compartment (Fig. 1C, p=0.005) and transitions to the light (Fig.1D, p=0.0002). The EPM test confirmed and strengthened the LDB outcomes. Indeed, after EIPV, the percentage of time spent in the maze open arms plummeted to almost zero (≅95% drop, p=0.002; Fig.1F). The same was evident for the percentage of entries in the open arms (pre vs. post =-90%, p<0.0001, Fig.1G). Notably, the freezing behavior increased markedly after EIPV (p=0.007, Fig.1H). Yet, locomotor activity was unaffected in these mice. Indeed, the total distance traveled by the female mice and average speed were unchanged after EIPV (Fig.1I-J). Finally, circulating corticosterone, a marker widely used in stress paradigms, was significantly increased after 21 days of EIPV (p=0.024, Fig.1K). Thus, the EIPV paradigm engendered a forthright anxiety-like behavior framed into a persistent general stress context, all typical features of human IPV.

### EIPV triggers apoptosis and halts neurogenesis at the hippocampal DG level

The hippocampus, particularly the dentate gyrus (DG), is a primary hub for mood control, memory consolidation, and spatial exploration^5,13,14^. Of relevance, alterations at this level are manifest in experimental and human forms of acute or chronic stress^6,15,16^. Yet, no studies have evaluated the impact of reiterated IPV-like conditions on the hippocampus, specifically the DG. To fill this gap, we first assessed the amount of apoptosis as a primary marker of cell damage. TUNEL assay revealed that cell death extent in the DG doubled after EIPV, as indicated by the increased number of TUNEL-positive nuclei spread throughout this structure, i.e., in the granular, subgranular, and hilus zones (Fig.2B, C) (control vs. EIPV; p=0.008). Next, we quantified hippocampal neurogenesis. To this end, we injected BrdU (75 mg/kg x 3 times) intraperitoneally to control (i.e., no EIPV entering) mice and those randomized to undergo the full EIPV protocol. After 21 days of EIPV, the number of BrdU+ cells, colocalized with doublecortin (DCX) positive nuclei, was markedly reduced in the DG (Fig.2D-E; control vs. EIPV p=0.008). DCX is a protein associated with neuronal differentiation and functionally linked to cell migration^17^. DCX is only expressed in cells contributing to adult neurogenesis in the DG, and previous studies^18^ show that DCX+ cells can be subtyped according to the neurogenic process stage, from immature through proliferative to postmitotic (Fig.2F). Hence, we next analyzed the number and percentage of DCX+ cells in the postmitotic phase^18^. We found markedly lower DCX+ postmitotic cells in the DG of EIPV-inflicted subjects than controls (Fig.2G-J, p=0.039, p=0.008). When assessing the distance from the soma of the DCX dendrites as an index of arborization and differentiation, we found it sizably reduced in the EIPV group (Fig.2K, p=0.008). These data reveal that the EIPV paradigm can induce cell death and impair the neurogenic process in the hippocampal DG.

**Figure 2.**
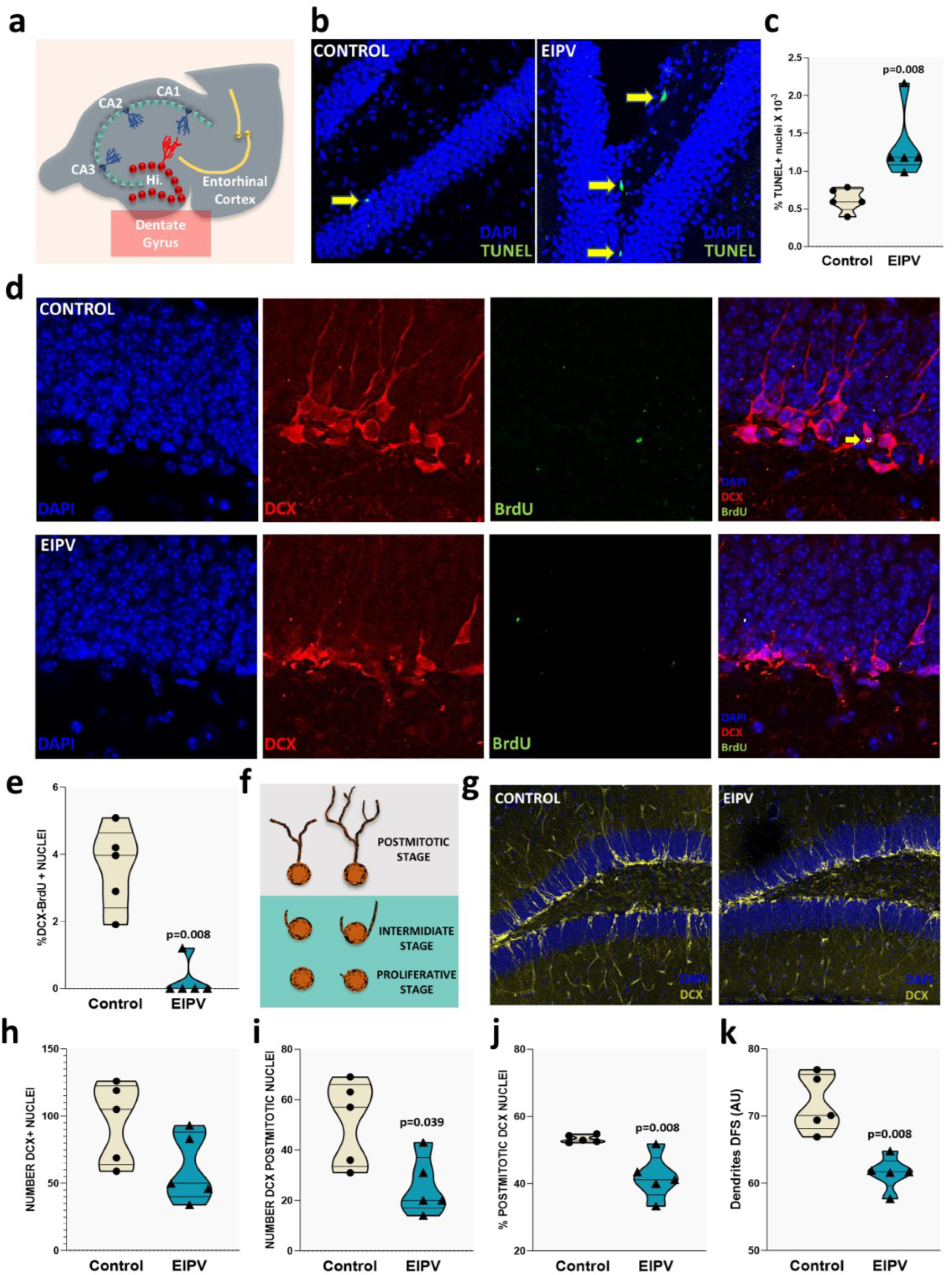
EIPV triggers apoptosis while impairing neurogenesis in the DG. **a**, Scheme of the hippocampal formation morphology. **b,** Representative images of IF with TUNEL assay in the DG of Controls and EIPV-inflicted mice. **c,** TUNEL positive nuclei quantification, control Vs. EIPV, n=5, p=0.08. **d,** Representative images of IF staining for DAPI, BrdU, and DCX in the DG of Controls and EIPV-inflicted mice. **e,** BrdU/DCX positive nuclei quantification, Control Vs. EIPV, n=5, p=0.08. **f,** Representative scheme of DCX positive cell differentiation. **h,** DCX positive nuclei quantification, Control Vs. EIPV, n=5. **i,** DCX positive nuclei (classified as postmitotic) quantification, Control Vs. EIPV, n=5, p=0.039. **j,** Percentage of DCX postimitotic positive nuclei (on total DCX positive nuclei), Control Vs. EIPV, n=5, p=0.008. **k,** Average measure of dendrites length of DCX postmitotic positive nuclei, Control Vs. EIPV, n=5, p=0.008. For all the results shown in the figure, it has been performed a Mann-Whitney U test between values from controls and EIPV-inflicted mice.

### EIPV impairs ERβ-dependent signaling in the hippocampus/DG of female mice

Estrogens are involved in the brain’s sexual differentiation and several neuronal events, such as survival and synaptic plasticity^19,20^. Moreover, estradiol (E2) governs mood balance and pain regulation, mainly through its specific receptors ERα and ERβ^21^. Since no previous studies have tested whether estrogens and their receptors are modified in human or experimental IPV, we first measured the circulating and whole brain levels of E2 before and after EIPV. Differently from our initial hypothesis, we found that, after EIPV, both systemic and central E2 levels were above pre-EIPV values (Fig.3A,B). In small and large mammals, ERα and ERβ are expressed all across the brain. Yet, ERβ expression is particularly prominent in the hippocampus, especially in the DG^22^ (Fig.3C). Accordingly, we observed a substantial drop in ERβ expression and unchanged levels of ERα at this level (Fig.3D-F). When zooming on the DG via IF, we noted a 70% decline in ERβ expression in EIPV female mice (p<0.0001, Fig.3G, H). We interpret these data to indicate that EIPV hinders ERβs and dependent signaling without overtly affecting circulating E2 content.

**Figure 3.**
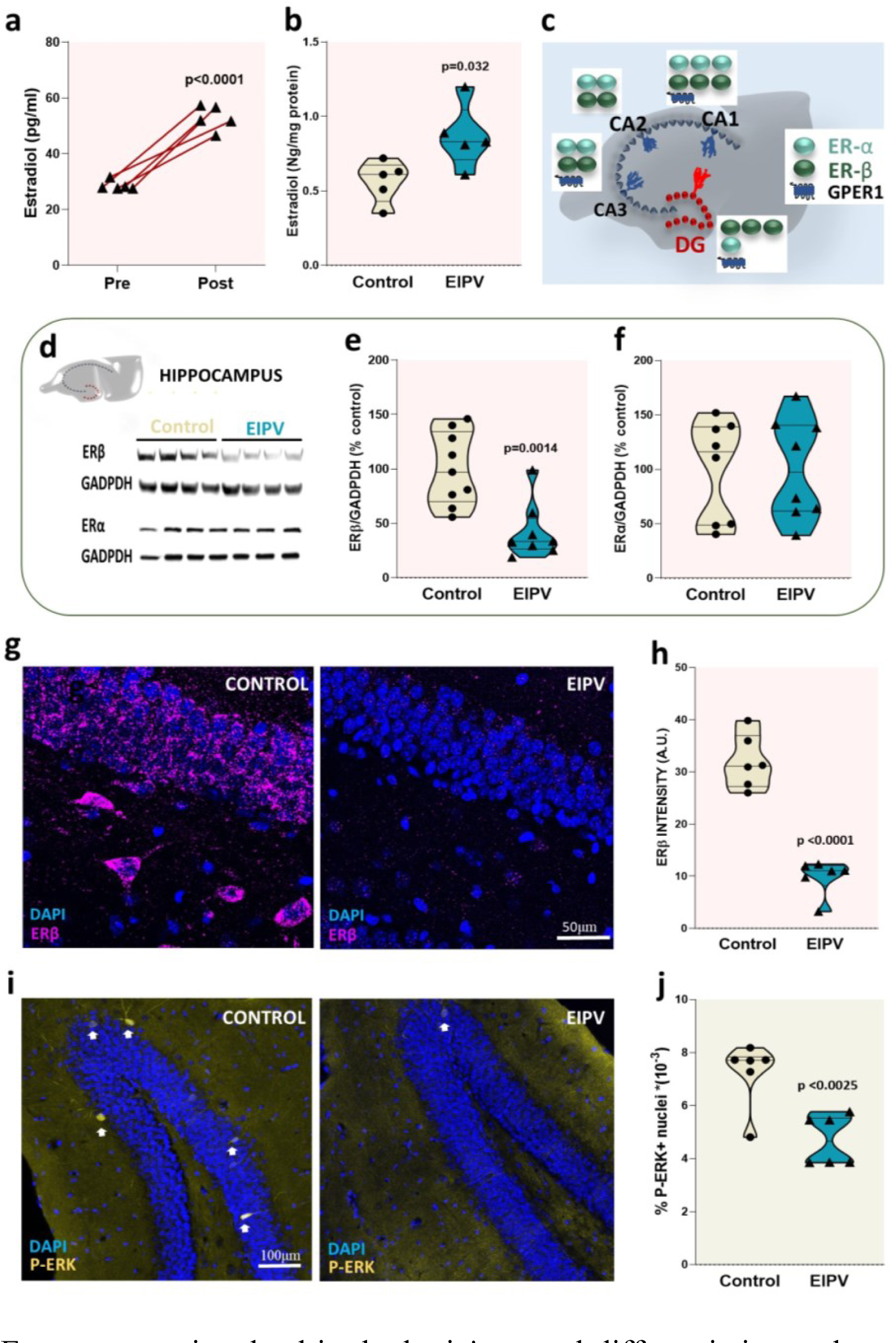
EIPV jeopardizes ERβ/BDNF signaling in the female hippocampus. **a**, Circulating 17-beta-estradiol measured by ELISA at Baseline and after 21 days of EIPV, n=5, p<0.0001. **b,** 17-beta-estradiol measured by ELISA in lysates of whole brain from controls and EIPV-exacted mice, n=5, p=0.022. **c,** Scheme of estrogen receptors expression in different hippocampal areas. **d-f,** Western blot for ERα, ERβ in hippocampus of controls and EIPV-imposed mice, (d) representative immunoblotting in the hippocampus for ERα, ERβ and GAPDH, (e) quantification of ERβ normalized on GADPDH content), control vs. EIPV, n=8, p=0.0014, (f) quantification of ERα (normalized on GADPDH content), control vs. EIPV, n=8. **g,** Representative images of IF staining for ERβ in the DG, control, and EIPV-inflicted mouse. **h,** Quantification of IF staining for ERβ in the DG, control vs. EIPV, n=6, p<0.0001. **i,** Representative images of IF staining for P-ERK in the DG, control, and EIPV-exacted mouse. **j,** Quantification of IF staining for P-ERK in the DG, control vs. EIPV, n=6, p=0.0025.

### EIPV fuels stress hormones-related signaling while downsizing the BDNF/TrkB axis in the DG

Estrogens can directly induce BDNF expression via ERs interaction, especially ERβ/ERK signaling^7,23^, while glucocorticoids calibrate BDNF expression in a dose- and condition-dependent manner^24^.

Major depressive disorders typically couple with BDNF downregulation and a glucocorticoid surge^25^, while chronic exposure to dexamethasone via glucocorticoid receptor (GR) activation can suppress BDNF expression in mouse hippocampal cells^26^. On these grounds, we determined whether a lack of ERβ and/or TrkB signaling accounts for reduced ERK activation, i.e., phosphorylation, leading, in turn, to BDNF depletion in the hippocampus. EIPV markedly dropped ERK phosphorylation levels in the hippocampal DG (p=0.0025; Fig.3I-J).

Estrogens limit the glucocorticoid response and its sequelae, such as inflammation^27^. At the hippocampal level, estrogen, glucocorticoids, and BDNF/TrkB entertain a complex relationship to govern adaptative responses to stress conditions^28^. Systemic levels of corticosterone are markedly elevated after EIPV (Fig.1K). Hence, we interrogated whether this surge permeates the brain, too. We found that EIPV boosted corticosterone levels in the whole brain (Fig.A, p=0.006). Corticosteroid hormones can bind mineralocorticoid (MR) and glucocorticoid (GR) receptors. Hence, we next interrogated whether a heightened GC activity in the DG paralleled brain corticosterone elevation. Accordingly, we evaluated the extent of GR phosphorylation level in DG via IF. Consistent with our original hypothesis, EIPV treatment induced a prominent rise in the GR phosphorylation magnitude in the DG (Fig.4B, C, p=0.0079).

**Figure 4.**
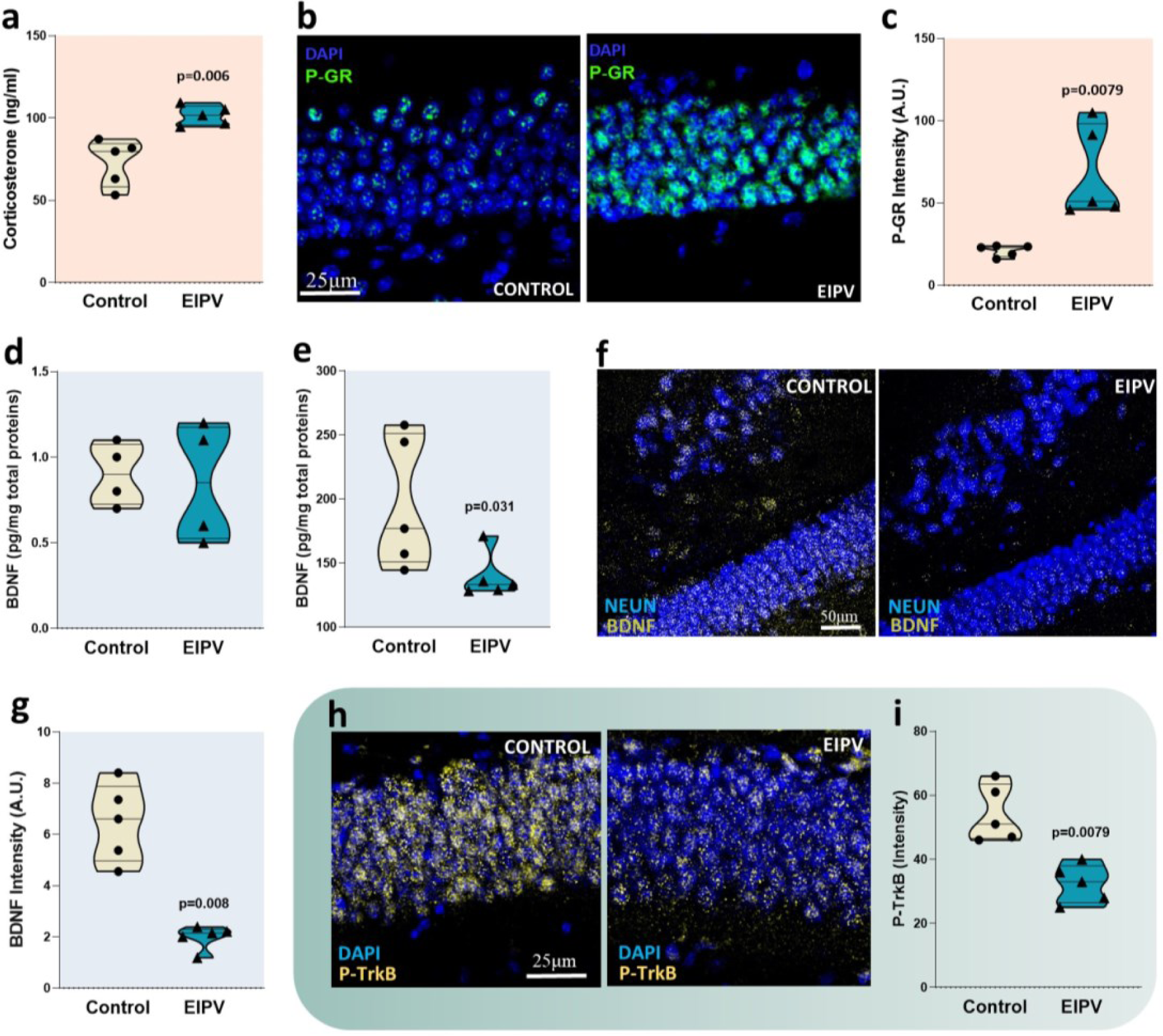
EIPV alters the balance between stress hormones and BDNF/TrkB signaling in the DG. **a**, Whole brain corticosterone levels measured by ELISA assay, control vs. EIPV, n=5, p=0.006. **b,** Representative images of IF staining for P-GR in the DG. **c**, Quantification of IP staining for P-GR in the DG, control vs. EIPV, n=5, p=0.0079. **d**, Whole BDNF content via ELISA assay, control vs. EIPV. **e,** Hippocampal BDNF measured by ELISA assay, control vs. EIPV, n=5, p=0.031. **f**, Representative images of IF staining for BDNF in the DG, control, and EIPV-imposed mice. **g,** Quantification of IF staining for BDNF in the DG, control vs. EIPV, n=5, p=0.008. **h,** Representative images of IF staining for P-TrkB in the DG, control, and EIPV-mice. **i,** Quantification of IF staining for P-TrkB in the DG, control vs. EIPV, n=5, p=0.0079. For all the results shown in the figure, a Mann-Whitney U test was performed between controls and EIPV-imposed mice values.

Next, we tested whether these events prompted local impoverishment, i.e., hippocampal/DG, BDNF pools. When examining BDNF levels in the whole brain via ELISA, we first found no significant differences between control and EIPV-inflicted females (Fig.4D). However, when zooming on the hippocampus, we noticed a noticeable BDNF depletion (Fig.4E, p=0.031). Consistent with the above-reported findings, this decline was even more pronounced in the DG (−70%, Fig.4F, G, p=0.008). In unison, we also detected a significant decrease in TrkB phosphorylation (Fig.4H, I, p=0.0079) in EIPV-inflicted mice. Thus, EIPV depletes hippocampal BDNF pools, especially in the DG. We can ascribe this decline to a lack of ERβ/ERK and/or TrkB agonistic signaling, on the one hand, and/or enhanced suppressive action exerted by chronic GR activation, on the other.

### Deleting ERβ resembles EIPV-ignited behavioral and molecular signatures

To cement the view that ERβ is causally involved in the abovementioned post-EIPV behavioral alterations, we extended our studies to ERβ KO mice (Fig.5A,B), evaluating them at baseline and after EIPV. At baseline, EβR KO mice already displayed a constitutive anxiety-like behavior, as evidenced by the EPM test’s main parameters, characterized by a significant decrease of time and percentage of entries in the open arms of the maze (Fig.5C-E). In unison, detecting BDNF via IF at the DG level revealed a drastic drop in basal expression of this neurotrophin in ERβ KO vs. WT mice (Fig.1A,B Sup.Mat.). We then subjected WT and ERβ KO mice to the full EIPV protocol, evaluating survival via the Kaplan-Meir curve. All WT mice survived the entire EIPV procedure. Conversely, four out of nine ERβ KO mice died prematurely during it (Fig.5I). Finally, the E2 circulating levels rose significantly after EIPV in WT mice, as shown in Figure 3A. Conversely, ERβ KO mice already had substantially higher systemic E2 at baseline than WT. In the former, EIPV still heightened E2 circulating levels, which did not outweigh those found in post-EIPV WT mice (Fig.5H). These data consolidate the view that alterations in the ERβ signaling underscore, at least, some of the EIPV-triggered molecular and behavioral features.

**Figure 5.**
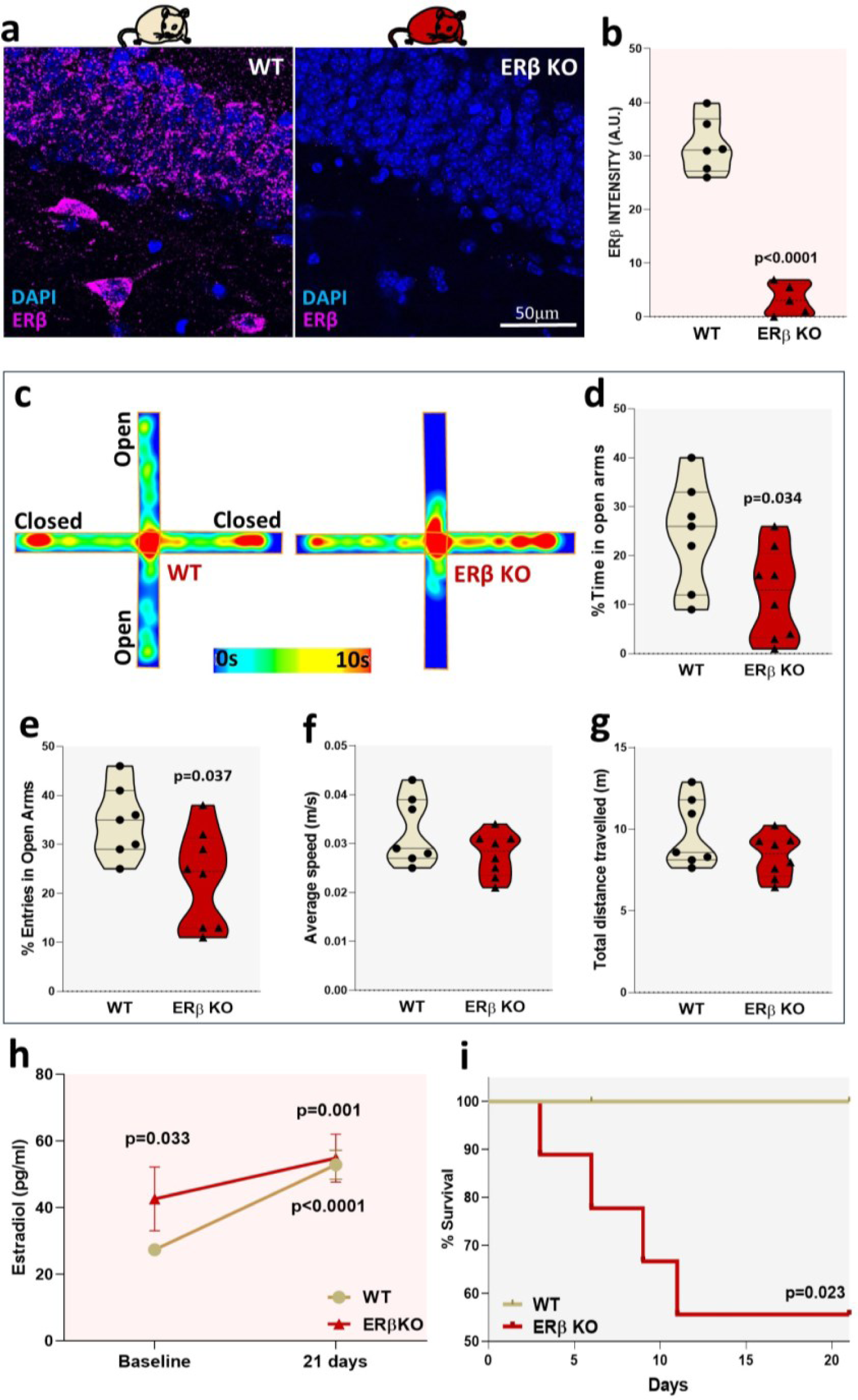
Genetic deletion of ERβ mimics EIPV-ignited behavioral and molecular alterations. **a**, Representative images of IF staining for ERβ in the DG, WT, and ERβ KO mice. **b,** Quantification of IF staining for ERβ in the DG, control WTs vs. ERβ KO mice, n=6/5, p<0.0001. **cg,** Elevated Plus Maze Test, (c) Heat plots of time spent in different parts of the labyrinth for WT (left panel) and ERβ KO mice (right panel), (d) Percentage of time spent in the open arms, WT vs. ERβ KO mice, n=7/8, p=0.034, (e) Percentage of entries in the open arms, WT vs. ERβ KO mice, n=7-8, p=0.037, (f) Average speed of the mouse movements, WT vs. ERβ KO mice, n=7/8 (g) Total distance traveled by WT and ERβ KO mice, n=7/8, **h,** Circulating 17-beta-estradiol measured by ELISA at baseline and after 21 days EIPV in WT and ERβ KO mice, n=5, baseline WT vs. baseline ERβ KO mice p=0.033, WT baseline vs. 21 days EIPV, p<0.0001, ERβ KO mice baseline vs. 21 days EIPV, p=0.001. **i,** Survival Kaplan-Meir curves of WT and ERβ KO mice during EIPV procedure, n=9, p=0.023.

**Figure 6.**
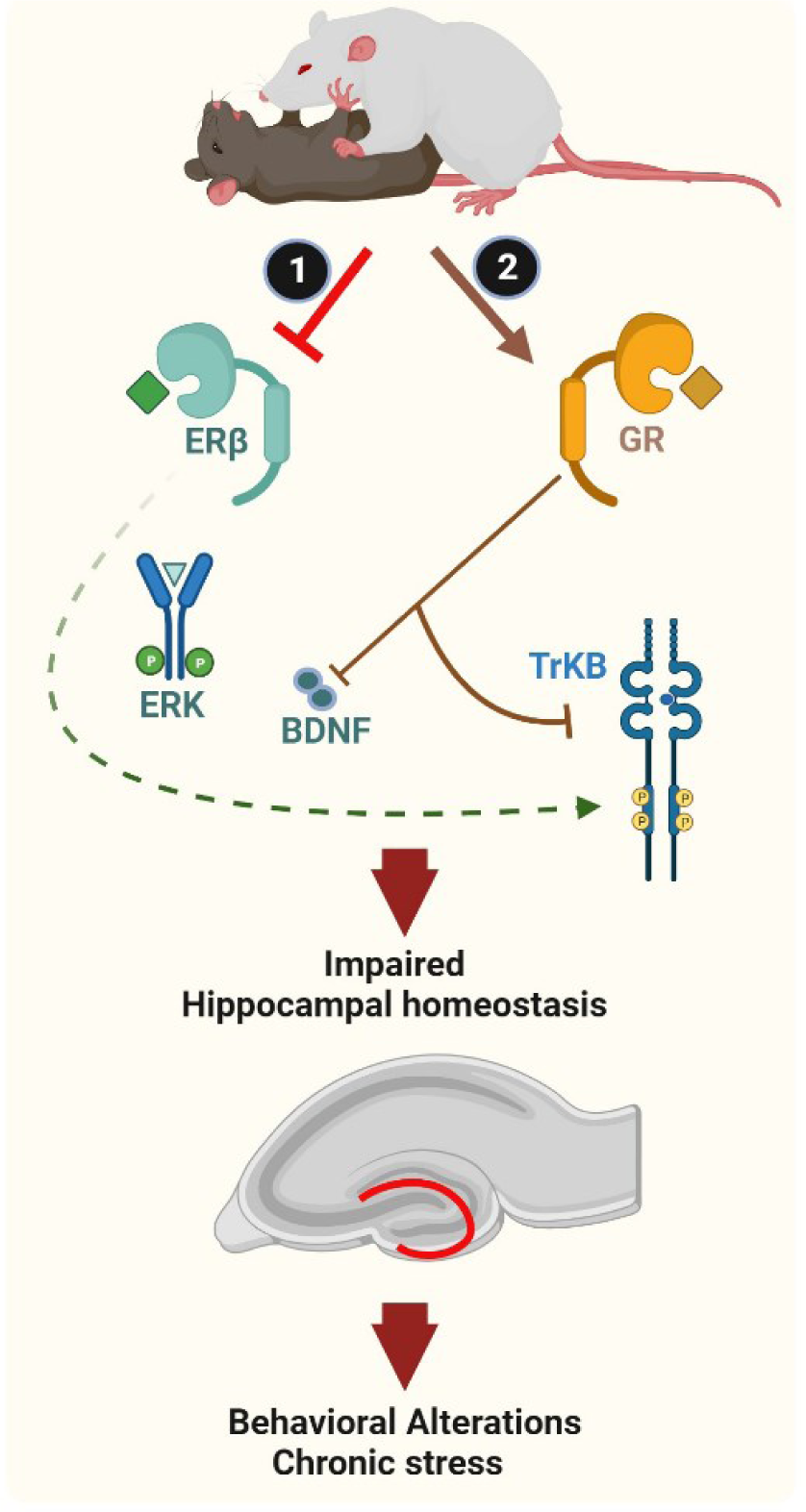
Synopsis of the main findings. Reiterated EIPV enacted by an aggressive CD1 male mouse on a C57BL6/j female mouse down-sizes ERβ receptor density in the female hippocampus (DG), lowering ERK-dependent BDNF generation, thus TrkB phosphorylation/activation. In parallel, EIPV triggers a corticosterone surge coupled to GR activation that contributes to dampen BDNF/TrkB signaling. Pathways 1 and 2 conjure up to impair neurogenesis in the female hippocampus, accounting for EIPV-driven behavioral abnormalities and chronic stress.

## Discussion

In 2022, the EU Agency for Fundamental Rights randomly performed interviews among the 28 countries of the UNION with 42,002 women aged 18-74, 51.7% of whom reported being violence victims in their lifetime^29^. On the heels of this evidence, getting a deeper grasp of the pathophysiological IPV sequelae becomes increasingly pressing. Here, we show that experimental IPV: a) propels anxiety-like behavior in female mice; b) fuels apoptosis and impairs neurogenesis at the hippocampal level; c) jeopardizes ERβ-dependent protective signaling, increasing stress hormone-related action while downsizing the prosurvival/connectivity-enhancing properties of BDNF/TrkB signaling in the DG.

We have adopted and modified a method for mimicking some prominent features of human IPV, such as reiterated physical and psychological harm for male-to-female. We focused on the hippocampus, especially the dentate gyrus (DG) given its vulnerability to social stress conditions^12,13^, and its prominent ERα and ERβ expression, eminent in the DG^22^.

After continuing physical attacks/sensorial intimidation by an aggressive male mouse, female mice develop anxiety-like behavior and chronic stress, two main features of human IPV. Although not the sole mechanism at play, our data indicate male violence curtails ERβ receptor density in the female’s hippocampus, particularly in the DG. At this level, the lack of this protective signaling prompts neuronal cell death, deficient dendritic arborization, and impaired neurogenesis. This adverse remodeling owes, at least in part, to the defective estrogen-evoked activation of master regulators of cell behavior, such as ERK and its dependent prosurvival signaling, including, but not limited to, BDNF expression.

Of note, in humans and other species, the link between hippocampal homeostasis (essential for affect regulation, stress response, memory, and spatial orientation) has been primarily studied in the context of psychopathology and aging in males^30,31^. Here, we extend it to reiterated physical violence suffered by the female at the hands of a male conspecific.

Although ERα and GPER1’s role in EIPV hippocampal pathophysiology warrants further investigation, our data suggest a primary involvement of the ERβ isoform, which is also the most expressed in this brain area^22^, fitting nicely with consolidated evidence that ERα and ERβ exert opposing actions during stress responses^32^. Accordingly, Krezel and colleagues reported a heightened anxiogenic behavior in ERβ KO but not ERα KO mice^33^. Equally relevant, the highly potent ERβ agonist diarylpropionitrile, DPN, increases open field entries and time spent in the open arms in the elevated plus maze in WT but not in ERβ KO mice^32^. While validating this notion, our current data also provide unprecedented evidence that stress conditions, such as EIPV infliction, directly hinder ERβ density. In our model, the elevated (and likely compensatory) systemic E2 levels can favor binding with the α isoform, whose expression, at least at the hippocampal level, remains unchanged, paradoxically exerting a negative effect on the pathways governed by the β isoform. However, as alternative or additive mechanistic route underscoring ERβ downregulation, we could also entertain the methylation of the ERβ promoter (5’ untranslated region), increasingly recognized as a primary modality for regulating this ER isoform extent^34^. From a behavioral point of view, ERβ modulation is critical for serotonin and dopamine neurotransmission^35^. Thus, the current lack of ERβ stimulation fuels the anxiety-like behavior found in EIPV-inflicted WT mice and exacerbated in ERβ KO ones.

At the hippocampal level, estrogen, corticosteroids, and BDNF influence each other levels^28,36^, thus the adaptative response to stress conditions^26^. Under the current scenario, we speculate that lack of ERβ, therefore unopposed corticosteroid receptor stimulation, contributes to the overall hippocampal structural and functional impairment by affecting, among other effects, TrkB activation status. The current evidence of a reduced extent of TrkB phosphorylation after EIPV grants support this contention.

We can envision future, in-depth studies employing *in vivo* imaging and electrophysiology approaches to more longitudinally track down hippocampal and brain dynamic perturbations. Nevertheless, our intent here was to provide an initial mechanistic template, i.e., EIPV disruption of the ERβ/ERK/BDNF axis, on which to mold additional evaluations of other brain regions equally relevant for behavioral homeostasis, such as frontal cortices, insula, amygdala, hypothalamus.

Our study demonstrates that recurrent physical/sensorial violence directly impacts the female hippocampus, accounting for primary shifts in behavioral control, such as the onset/maintenance of anxiety and chronic stress. At the same time, from a therapeutic perspective, our current data suggest that, in IPV victims, preserving ERβ integrity should be at the core of the therapeutic attempts. This proposition has additional significant clinical implications. Indeed, if generalized, the loss of ERβ signaling could favor the progression of many forms of cancer in women. Unfortunately, cancer diagnoses are increasing in IPV-victimized women^37^. Moreover, studies suggest ERβ activation may be a promising avenue for reducing menopause-related hot flashes, and memory dysfunction, and also to lessen the risk of neurodegenerative disorders, such as Alzheimer’s disease^38^.

Women victims of EIPV show a plethora of pathological signs and symptoms spread throughout almost all body areas. Often, these conditions are treated separately and symptomatically. By grouping them in one common pathogenic mechanism, i.e., disrupted ERβ/BDNF signaling, we offer new potential therapeutic avenues, such as selective ERβ agonists derived from natural compounds ^39,40^, to taper off the acute and chronic central repercussions of reiterated IPV.

## Methods Behavioral tests

### Experimental Intimate Partner Violence

We employed a recently validated protocol to induce social defeat in female mice^4^ [in turn borrowed from the resident-intruder paradigm, widely used to research human psychosocial stress through animal models] to reproduce some salient conditions of human IPV (Fig.1A). In detail, we sprayed a 60 µl aliquot of urine collected from a CD1 male mouse on the genital area of a 12 weeks old female C57BL6/J mouse (Fig.1A, first step). Then, we placed the female mouse inside a cage containing a resident aggressive CD1 male mouse. We selected this dominant male mouse according to the criteria of the classical resident-intruder paradigm^10^; only the most aggressive males have been employed in our study (Fig.1A, step 2). This intervention triggered a violent reaction by the CD1 resident mouse, manifested in numerous attacks and bites during an interaction lasting 10 minutes (step 3). After this step, the two animals were maintained in the same cage for 24 hrs. but physically separated via a perforated transparent plexiglass barrier, thus allowing a continued sensorial connection, thus a psychological threat for the female (step 4). This procedure was repeated for 21 days to induce stress/social defeat. During the 10 min of interaction the main behavioral parameters linked to aggression and social defeat were recorded (i.e., the frequencies, durations, latencies of the attacks). To prevent animals’ injury, the procedure was interrupted in case of physical damages.

### Light-Dark Box

The light-dark box (LDB) measures anxiety-like behavior in female mice^11^. This test evaluates the innate aversion to open illuminated spaces and the spontaneous exploratory activity of mice. The test apparatus consists of a box separated into a smaller (one-third) dark chamber and a larger (two-thirds) brightly illuminated chamber. Mice were allowed to move freely between the two areas for 5 min while a camera recorded their behavior. The total time spent in the lighted compartment is an index useful to evaluate anxiety-like behavior. At the same time, the number of transitions between the two chambers is an index of exploratory behavior^41^.

### Elevated Plus Maze

The Elevated Plus Maze (EPM) evaluates anxiety-like behavior and locomotor activity in rodents^11^. The test equipment consists of four arm labyrinth, two protected by lateral walls (closed arms, 30 cm × 36 5 cm, × 15 cm) and the other two exposed (open arms, 30 cm × 5 cm, 350 lx). EPM exploits the rodent’s conflict between aversion to open spaces (i.e., hiding in closed arms of a labyrinth) and instinct to explore new environments (i.e., exploration of open arms of the same labyrinth). The experimental sessions have been videotaped by a camera placed above the apparatus. The analysis of the behavioral parameters has been performed employing ANY-maze software.

## Biochemical Assays

### Immunofluorescence

Animals were sacrificed with an overdose of ketamine–xylazine solution (K, 100 mg/kg; X, 10 mg/kg) and transcardially perfused with PBS 1X, followed by paraformaldehyde (PFA) 4% solution. The brains were removed from the skull, kept in the fixative (PFA 4%) for 24 hrs., and transferred into PBS 1X. A sliding microtome cut 50 μm coronal hippocampus sections in an anteroposterior direction (distance from Bregma, rostral to caudal from −1.94 mm to −3.64 mm). For each immunostaining (except for the TUNEL assay), free-floating sections were pretreated with 70% formic acid for 20 minutes at room temperature. Then the slices were blocked for 1 hr. at room temperature (RT) with 2% BSA, incubated overnight at 4 °C with primary antibodies (ERβ 1:200, Invitrogen; P-ERK 1:100 CellSignaling; BDNF 1:200, Alomone Labs; P-GR 1:250, Cell Signaling, P-TrkB 1:250, EMD Millipore) and finally incubated with secondary antibodies (1:500, Alexa Fluor, Invitrogen) for 1 hr. at room temperature.

### TUNEL assay

TUNEL assay was used to detect hippocampal apoptosis (In Situ cell death detection kit; Sigma, Saint Louis, MO, USA), as described previously^15^. Percent apoptotic hippocampal cells were calculated via the number of TUNEL-positive nuclei over the total number of hippocampal nuclei per field. At a magnification of 20×, five confocal (Leica SP5) images were taken per mouse hippocampus on different z sections covering 20 μm.

### Adult neurogenesis

To evaluate adult hippocampal neurogenesis, BrdU (Sigma-Aldrich) was administered intraperitoneally at 75 mg/kg body weight (divided into three different injections every 2 hrs.) the day before starting the EIPV procedure and evaluated at the end of the 21 days by IF staining. The number of doublecortin (DCX) (Abcam) and BrdU (Cell-Signaling) positive cells was estimated in serial coronal sections covering the complete rostrocaudal extension of the dentate gyrus (DG). At a magnification of 20×, five confocal (Leica SP5) images were taken per mouse hippocampus on different z sections covering 20 μm). DCX-expressing cells were classified into six distinct groups according to their morphology as previously described^18^ (Fig. 2F). For each DCX-expressing cell, classified as E and F, ImageJ measured the most extended dendrite length.

### Protein expression quantification

For ERβ, BDNF, and P-GR expression levels, the following method of quantification was employed: the mean fluorescence intensity within the granular layer and hilus (DG) were quantified using Fiji software (NIH) and with background (same fluorescent signal belonging to an area selected outside the granular layer and free of DAPI positive nuclei) subtracted from each image, according to previous studies^42^. Images were acquired via a Leica SP5 confocal microscope.

### Enzyme-linked immune assay

Tissues from the hippocampus and whole brain were dissected and frozen in isopentane. Tissues were homogenized in 2–3ml of lysis buffer (100mM PIPES (pH 7), 500mM NaCl, 0·2% Triton X-100, 2mM EDTA, 200μM PMSF, and protease inhibitor cocktail (Sigma-Aldrich), followed by centrifugation for 10 min at 16000g at 4°C. BDNF protein levels were determined by Enzyme-Linked Immune Assay (ELISA) (BDNF Immuno Assay System, Promega; 17 beta Estradiol ELISA, Abcam). Lysates, after dilution, were loaded into 96-well plates. BDNF concentration (pg/ml) was normalized to total soluble protein previously measured through the BCA Protein Assay Kit (Pierce).

### Western Blot

Proteins were extracted from snap-frozen whole brain and hippocampal samples. We loaded forty micrograms of total protein lysate per lane on 4–20% precast polyacrylamide gel (Mini-PROTEAN® TGX™, Bio-Rad Laboratories) and blotted electrophoretically. The membrane, including samples was probed with specific antibodies (ERβ 1:500, Invitrogen; ERα 1:500, Invitrogen), and then re-probed for glyceraldehyde 3-phosphate dehydrogenase (GAPDH) with rabbit polyclonal antibody (dilution1:1000; EMD Millipore). Anti-GAPDH antibodies were used to verify the uniformity of protein loading. The protein bands were developed in a chemiluminescence substrate solution (Pierce SuperSignal Chemiluminescent substrate). Analysis of protein bands was performed using ImageJ software (National Institute of Health, USA). We checked the predicted molecular weight using Precision Plus Protein™ Dual Colour Standards (Bio-Rad Laboratories, Inc., Hercules).

### ERβ KO mice

For the studies shown in Figure 5, we employed 3-4 months old female ERβ KO mice from the Jackson Laboratory (stock #004745). These mice have been generated by inserting a neomycin resistance gene into exon 3 of the coding gene for ERβ by using homologous recombination in embryonic stem cells^43^.

### Statistics and Reproducibility

Results are presented as violin plots. All parametric data were analyzed by unpaired t-tests between the control and EIPV groups. We used the Mann-Whitney U Test for experiments where the sample size was less than six units per group. For the behavioral experiments and the circulating markers (longitudinal study), we employed paired t-tests between baseline and post-treatment. For experiments shown in Figure 5H (4 groups of animals), we used repeated measure ANOVA. All the statistical analyses were conducted through GraphPad PRISM 8.0.2.

## Acknowledgments

This study was funded by MSCA action (Marie Skłodowska-Curie Actions) PINK (to J.A.), EU VISGEN MSCA-RISE-2016, EU NEURAM FETOPEN-RIA-2014-2015 to (M.D.M), and NIH-NHLBI R01 HL136918 (to N.P.). Thanks to Dr. Anna Sanga for the digital scheme (Fig.1A).

## Author contributions

<u>Literature search:</u> Jacopo Agrimi, Marco Dal Maschio, Nazareno Paolocci. <u>Study design:</u> Jacopo Agrimi, Marco dal Maschio, Nazareno Paolocci. <u>Animal data collection:</u> Jacopo Agrimi, Lucia Bernardele, Naeem Sbaiti, Marta Canato, Ivan Marchionni. <u>Molecular and histological data</u> <u>collection:</u> Jacopo Agrimi, Lucia Bernardele, Naeem Sbaiti. <u>Data analysis:</u> Jacopo Agrimi, Lucia Bernardele, Naeem Sbaiti, Nazareno Paolocci. <u>Data interpretation:</u> Jacopo Agrimi, Beatrice Vignoli, Marco Canossa, Marco Dal Maschio, Nazareno Paolocci. <u>Drafting of the manuscript:</u> Jacopo Agrimi, Nazareno Paolocci. <u>Figures:</u> Jacopo Agrimi, Nazareno Paolocci. <u>Critical</u> <u>manuscript revision:</u> Jacopo Agrimi, Beatrice Vignoli, Ivan Marchionni, Marco Canossa, Nina Kaludercic, Claudia Lodovichi, Marco Dal Maschio, Nazareno Paolocci.

## Supplemental Materials

**Figure 1.**
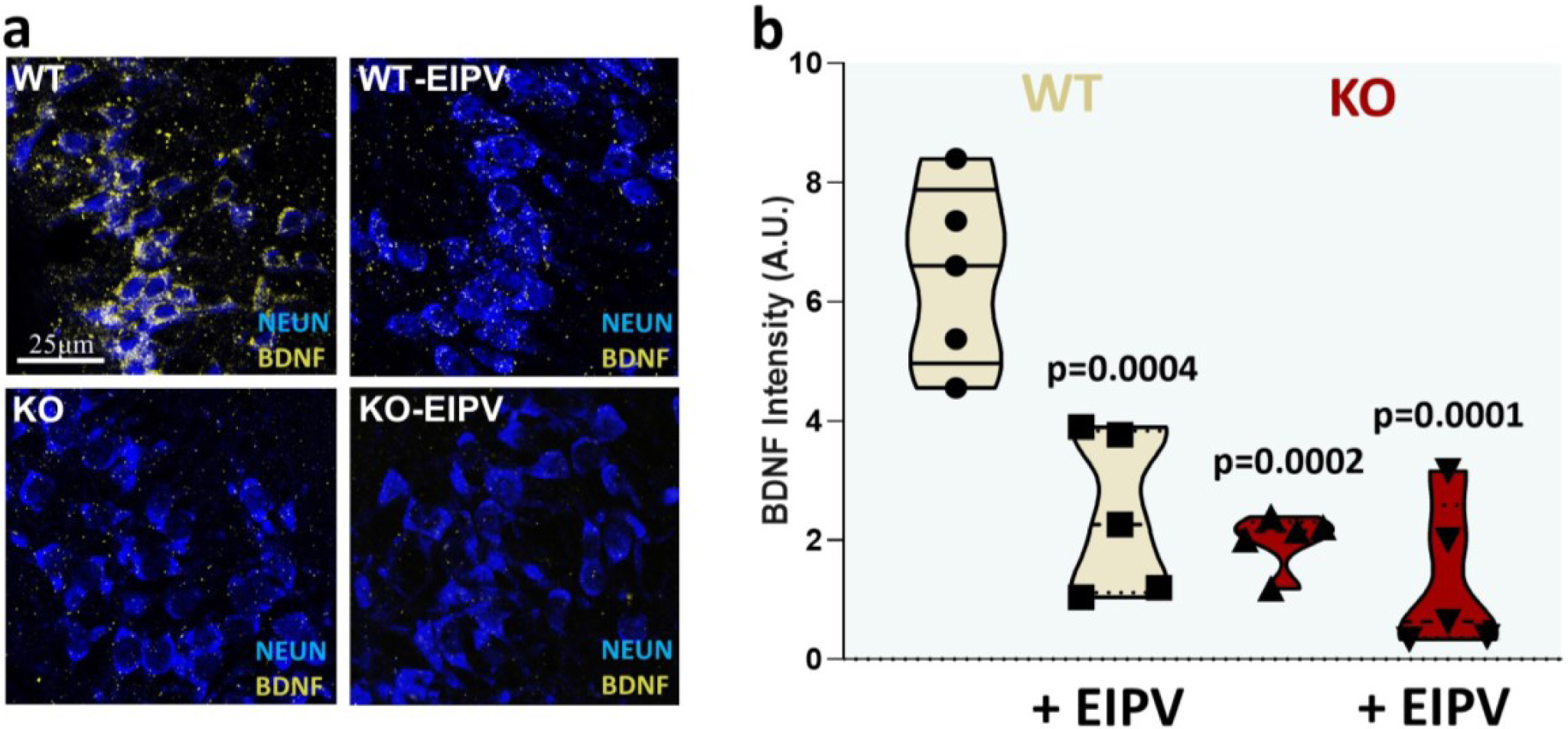
EIPV induces anxiety-like behavior: **a**, Experimental Intimate Partner Violence (EIPV), 60 μl aliquot of urine collected from a CD1 male mouse is sprayed on the genital area of a 12 weeks old female (first step); then, the female mouse is placed inside a cage containing a resident aggressive CD1 male mouse (step 2); this intervention triggers a violent reaction by the CD1 resident mouse during a phase lasting 10 minutes (step 3); after this step, the two animals are maintained in the same cage for 24 hrs. but physically separated via a perforated transparent plexiglass barrier, thus allowing a continued sensorial connection (step 4). The procedure is repeated for 21 days to induce stress/social defeat. **b**, Average quantification of the time spent under attack of the females subjected to EIPV. **c-d,** Light-dark box test, (c) Time spent by mice in the light compartment, Pre-EIPV Vs. 21 days EIPV, p=0.005, (d) Number of transitions from the dark to the light compartment; Pre-EIPV Vs. 21 days EIPV, p=0.002. **e-j,** Elevated Plus Maze Test, (e) Heat plot of time spent in different parts of the labyrinth at baseline (left) and after EIPV (right), (f) Percentage of time spent in the open arms; Pre-EIPV Vs. 21 days EIPV, p=0.002, (g) Percentage of entries in the open arms, Pre-EIPV Vs. 21 days EIPV, p<0.0001, (h) Time of freezing behavior, Pre-EIPV Vs. 21 days EIPV, p=0.0065, (i) Total distance traveled by the animals, Pre-EIPV Vs. 21 days EIPV, (j) Average speed of the animal movements, Pre-EIPV Vs. 21 days EIPV. **k,** Circulating corticosterone, Pre-EIPV Vs. 21 days EIPV, p=0.024. For all the results shown in the figure, a paired t-test was performed between values at baseline and after 21 days of EIPV, n=8.

## Notes

### Competing Interest Statement

The authors have declared no competing interest.

